# Genetic analysis using parent-progeny relationship for wood quality traits in Norway spruce (Picea abies (L.) Karst.)

**DOI:** 10.1101/293969

**Authors:** Linghua Zhou, Zhiqiang Chen, Sven-Olof Lindqvist, Lars Olsson, Thomas Grahn, Bo Karlsson, Harry X. Wu, María Rosario García Gil

## Abstract

Two-generations pedigree involving 524 plus trees and their open-pollinated (OP) progenies were jointly studied to estimate parent-progeny correlation and heritability. Three wood traits (wood density, MFA, and MOE) were determined by SilviS-can in one ramet per plus tree and 12 OP progenies. Three ramets per plus tree and 12 OP were also measured with two indirect methods, Pilodyn and Hitman. The overall correlation between OP-based breeding values and plus tree-based phenotypes was low to moderate for all traits. The correlations between the phenotypic values of the mother trees and the breeding values estimated on their half-sib pro-genies are low to moderate. Reasons for this may be experimental errors in progeny trials and lack of experimental design in archives, contributing to the parent and progeny correlation. The management practices in the archive may contribute more to such low correlation. Offspring progeny heritability estimates based on SilviScan measurements were higher than parent-offspring regression using one single ramet from the archive. Moreover, when three ramets were measured the parent-offspring regression heritability estimates were higher than those based solely on progeny data for the Pilodyn and Hitman on the standing trees. The standard error of the heritability estimates decreased with increasing progeny size.

## Introduction

Norway spruce [Picea abies (L.) Karst.] is one of the most important conifer species in Europe for both economic and ecological aspects [Hannrup et al., 2004].It is extensively forested for timber and paper production. Increasing volume production has traditionally been the main objective of the species breeding program, while more recently, different aspects related to wood quality have gained increasing attention [Mullin et al., 2011, Rosvall et al., 2011]. For mechanical properties of wood-based products, wood density, microfibril angle (MFA), and modulus of elasticity (MOE) are considered as the most important traits [Chen et al., 2015, Zobel and Jett, 1995].

The SilviScan technology was developed to measure the radial variation, i.e. from pith to bark of many trails from the same sample: solid-wood quality traits including wood density, MFA, MOE [Evans, 1999, Evans, 2006],as well as fiber traits, such as their cross-sectional dimensions and MFA [Evans, 1994].Its high resolution, accuracy and very high efficiency compared to corresponding laboratory methods, has contributed substantially to advancements in research and development within wood biology, forestry and optimal use of forest resources in softwoods [Lindstrm et al., 1998, Lundgren, 2004, Kostiainen et al., 2009, McLean et al., 2010, Piispanen et al., 2013, Fries et al., 2014], on hardwoods [Kostiainen et al., 2013, Lundqvist et al., 2017] and on modelling of trait variations [Wilhelms-son et al., 2002, Lundqvist S-O, 2005, Franceschini et al., 2012, Auty et al., 2014]. When data from thousands of samples are preferred, which is often the case in operational tree breeding, SilviScan is however seldom used. Then, rapid indirect methods may be better suited, if available and good enough. SilviScan is then typically used to produce benchmark data and for validation of the more rapid procedures. Examples of such faster and non-destructive methods (NDM) for solid wood traits are Pilodyn penetration and acoustic velocity [Chen et al., 2015, Kennedy et al., 2013,Vikram et al., 2011]. Pilodyn is an indirect non-destructive, low cost and easy-to-use instrument for measuring wood density. In Norway spruce and other conifer species, strong genetic correlations were observed between Pilodyn penetration and wood density measured with SilviScan [Chen et al., 2015, Cown, 1978, Desponts et al., 2017, Fukatsu et al., 2011, King, 1988, Sprague et al., 1983, Yanchuk and Kiss, 1993]. Further, acoustic velocity measured by Hitman ST300 (Christchurch, New Zealand) apparatus has shown an efficient indirect method for the estimation of MFA and it has already been used on many species, such as Scots Pine (Pinus sylvestris L.) [Hong et al., 2015], white spruce (Picea glauca [Moench.] Voss) [Lenz et al., 2013], and Norway spruce [Chen et al., 2015]. Models for estimation of wood stiffness from density and MFA have been developed for many species and were also implemented in early versions of SilviScan [Evans and Elic, 2001]. An analogous model using the proxy measurements of acoustic velocity and Pilodyn penetration on standing trees was shown to be efficient for the selection for wood stiffness in Norway spruce [Chen et al., 2015]. However, Pilodyn measures wood density only in the outermost annual rings, therefore, it has also been suggested that it may not be reliable for ranking of the whole tree in case where the correlation is low between the outermost rings and inner rings [Wessels et al., 2011] or if the diameter of tree is wide.

As part of the Norway spruce breeding program in Sweden, large numbers of superior genotypes (plus trees) are maintained in ex-situ grafted clonal archives. These serve as base breeding populations where crossings of selected parental genotypes are conducted with the purpose of generating cross-pollinated progenies for the next generation in the breeding cycle. There is an interest to estimate heritability using these clonal archive, particularly using the relationship between archived parent and their progenies, and clonal heritability (repeatability) for wood quality traits. Clonal archives are usually established using grafted scions from selected trees, without implementation of experimental design, and subject to management activities, such as pruning, topping and thinning at late ages. These operations may affect growth greatly but may have less impact on the genetic property of wood quality traits. This is because wood quality traits have higher heritability and little G by E [Chen et al., 2016, Hayatgheibi et al., 2017, Hong et al., 2014, Lenz et al., 2010, Wu et al., 2008]. Therefore, the parent-offspring relationship may still be used to estimate heritability for wood quality traits. The purpose of this study was to examine effectiveness of using parent-offspring to estimate heritability using the archived parent and progeny in field tests by:

1. Investigating the correlation and regression between the phenotypic values measured in clonal archives and the breeding values (BV) estimated in their half-sib progenies.
2. Comparing heritability estimates based on progeny only and based on parents-offspring regression and clonal repeatability for solid-wood quality traits.
3. Finally, we also investigated the effect of progeny and ramet number on heritability estimate.

## Materials and methods

### Plant material

The study was based on two generations of pedigree involving: (1) 524 mother plus-tree clones/genotypes from two different clonal archives in Ekebo and Mal-tesholm in southern Sweden. (2) Their corresponding 524 open-pollinated (half-sib) families consisting of at most six individuals for each of the two progeny trials. For the 524 mother trees, originally there were ca. 10 ramets graftted for each clone at the time of establishment, however, at the time of sampling the majority of the genotypes had only three ramets remaining. The trees were grafted on seedling rootstocks 28-30 years prior to sampling. As part of their maintenance, the clone archives had been thinned, and the trees topped. The two progeny trials were, S21F9021146 aka F1146 (Hreda, Eksj, Sweden) and S21F9021147 aka F1147 (Erikstorp, Tollarp, Sweden) were established in 1990. Increment cores had were sampled in 2010 and analysed with SilviScan at age 21-years old [Chen et al., 2014], whilst Pilodyn penetration depth and velocity were measured at ages 22 and 24, respectively [Chen et al., 2015].

### Measurements and data

The radial variations in wood density, MFA and MOE have been assessed in previous studies with SilviScan from increment cores of up to 12 progenies per half-sib family, followed by the calculation of area-weighted averages, representing the trait averages of all wood formed in the stem cross-sections at each cambial age. In this study, the corresponding data was generated also for the ramets from the archives. Below, widths of annual rings are used to identify outliers and DBH to compensate for differences in between tree competition on pre-treatments of data. For all other calculations, such as estimation of heritability and the comparison of methods, averages for stem cross-sections at different cambial ages (ring numbers) were used.

Data was also assessed through the use of the indirect proxies Pilodyn, Velocity and MOE(ind). The first two of these were measured on the standing trees with Pilodyn 6J Forest and Hitman ST300 instruments. MOE was estimated from these indirect measurements using the formula:

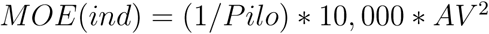

where Pilo is the Pilodyn penetration depth (mm) and AV is the velocity of the wave through the material (km/s), which has a high inverse correlation with MFA [Chen et al., 2014]. These data were previously presented in [Chen et al., 2014, Chen et al., 2015]. In this study, such data was generated on up to three ramets of each mother clone/genotype in the clonal archive. When more than one ramet was available, the average of each measurement was used for further Pearson correlation analysis. The evaluations were based on data for up to ring 16 to avoid problems of representability, as fewer trees were represented at the highest cambial ages.

### Data adjustments

The SilviScan data produced from the increment cores of the progeny trials when taken together and compared to those from the mothers, should be more informative about the family. They include data on up to 12 trees per family, whereas only one single ramet per each of the 524 mother tree genotypes was sampled for phenotyping with SilviScan at the clone archive level. Therefore, to identify outliers, we calculated for each family the 99% confidence intervals for the widths of rings with the same cambial age (ring number). The mother trees with more than seven rings of widths outside the 99% confidence interval of their progeny were considered outliers. When investigating the effect of removing outliers, these mother trees and their progenies were removed from the analysis.

For an overall comparison on how trees grown in the trials versus those in the archive had developed with age, averages for rings of the same number were calculated from SilviScan data across all progeny trees in the two trials and all mothers in the archive, respectively. These averages were plotted versus ring number (Figure 1), showing different characters of the developments of ring width and wood density between trials and archive. We assumed that this was caused by differences in between tree competition. To adjust the data for this, we constructed a competition index for the mother trees (Competition).

**Figure 1:**
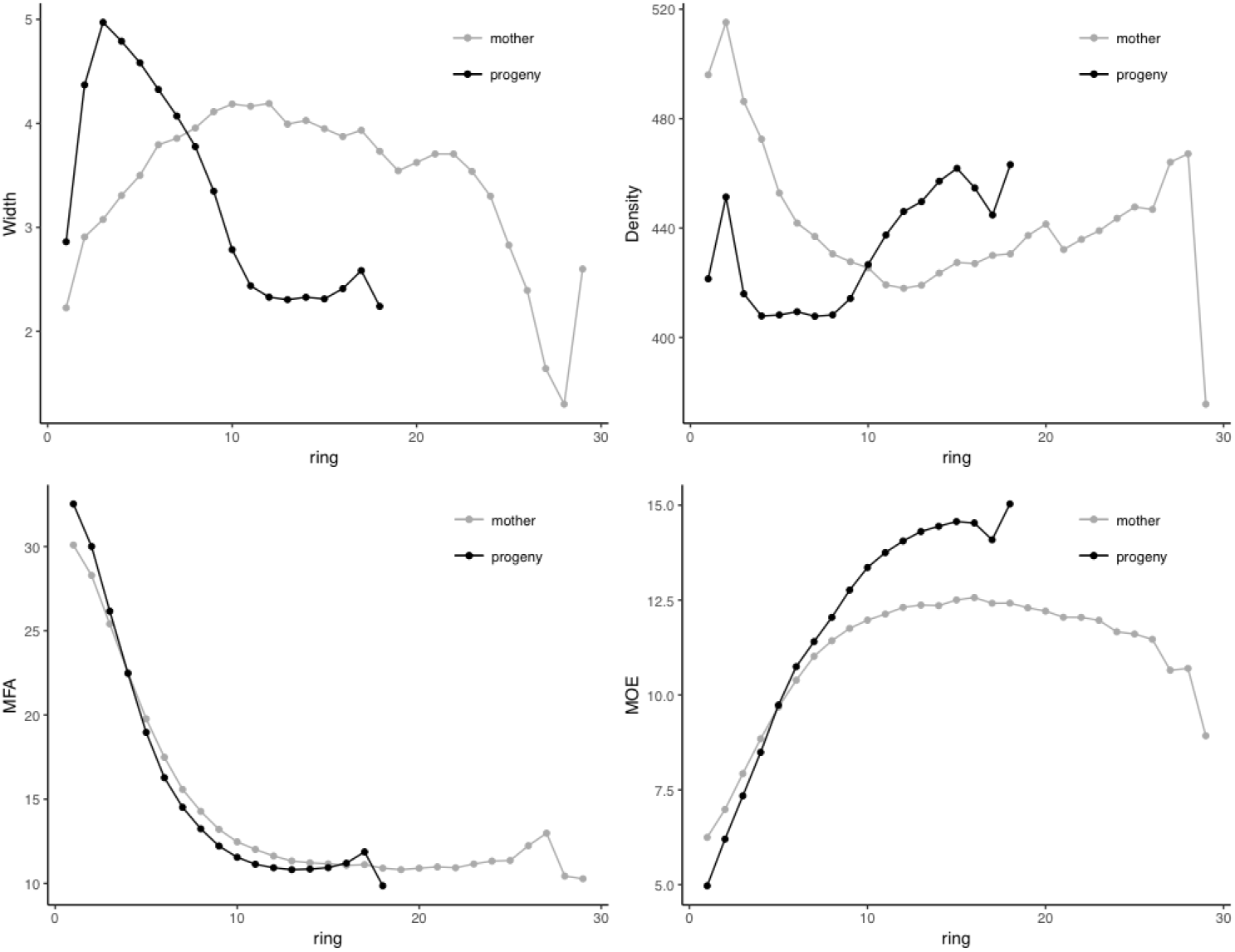
Mean values for all rings of same cambial age (ring number) across all progeny trees of all families, and across all mother trees in the archive, respectively, measured with SilviScan and plotted versus ring number for ring width, density, MFA, and MOE.

The original competition index is defined as follows [Hegyi, 1974]:

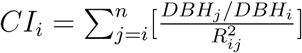

where *DBH*_*j*_ is the diameter at breast height at 1.3 m of jth neighbour tree, *DBH*_*i*_ is the diameter at breast height at 1.3 m of subject tree i, *R*_*ij*_ is the linear distance between ith subject tree and the jth neighbour tree. Since in this study, we only had the information of whether the DBH of the tree standing beside is larger or smaller than the subject tree, we set the ratio of DBH (nominator of this fraction) to be 0.5 if DBH of neighbour tree was smaller than subject tree; 1.5 if DBH of neighbour tree larger than subject tree and 1 if they had same DBH. Once the index was produced, it was used as a fixed effect for adjusting density, MFA and MOE.

Correlation and heritability estimates from un-adjusted and adjusted datasets are presented in Figure 2 and Figure 3, respectively. Data adjustment contributed little to the improvement of MOE estimates. For density, outlier removal resulted in the highest improvement, followed by adjustment for competition. For MFA, use of un-adjusted data provided the highest correlation and heritability, and outlier removal the lowest. Further heritability estimates were based only on data adjusted for competition.

**Figure 2:**
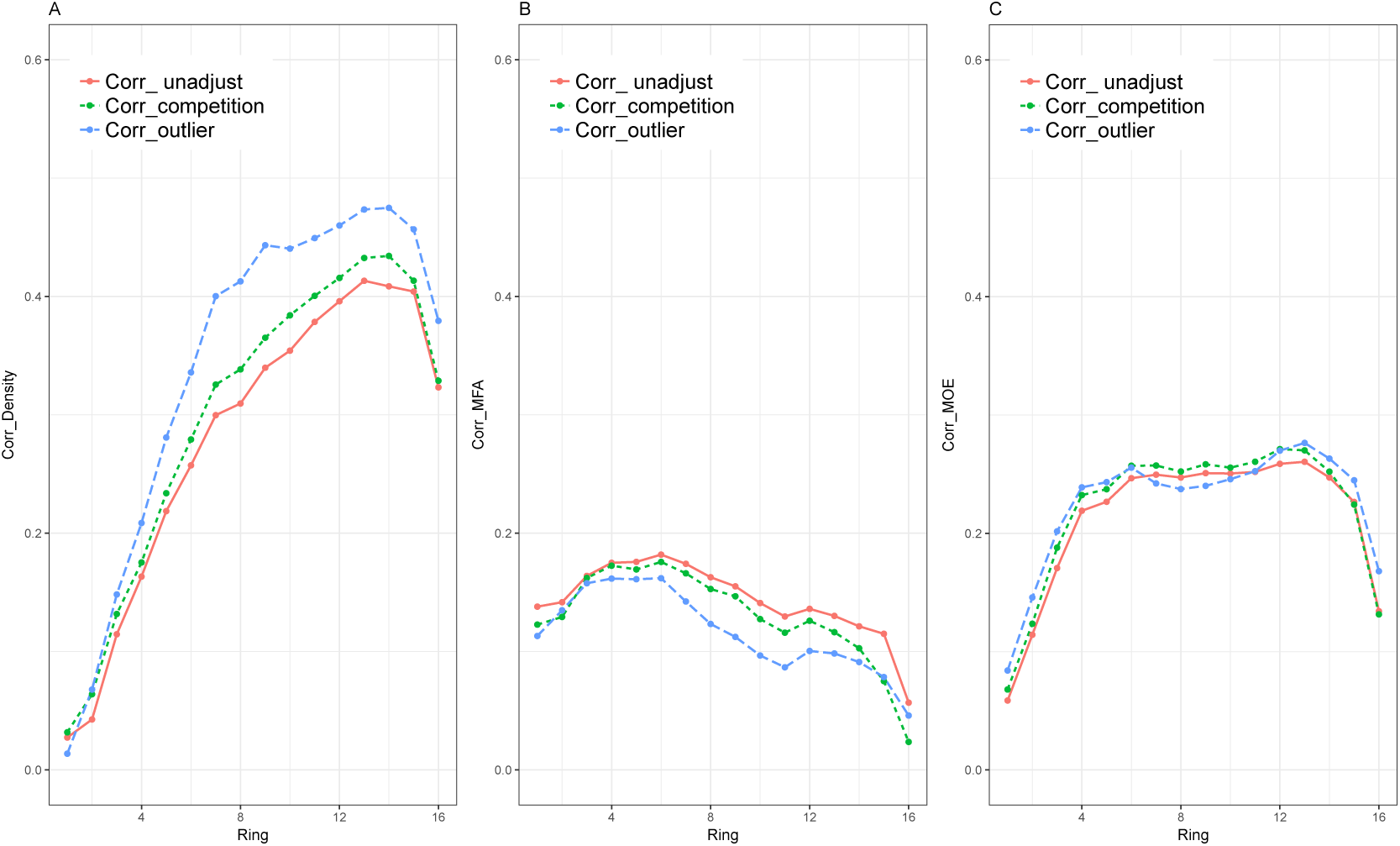
Correlations between BVs and phenotypic values of archive mother trees for area-weighted averages calculated from SilviScan data to represent the stem cross-sections at each cambial age (ring number), using three types of data: 1) un-adjusted, 2) after competition adjustment and 3) with outliers removed, for density (A), MFA (B) and MOE (C).

**Figure 3:**
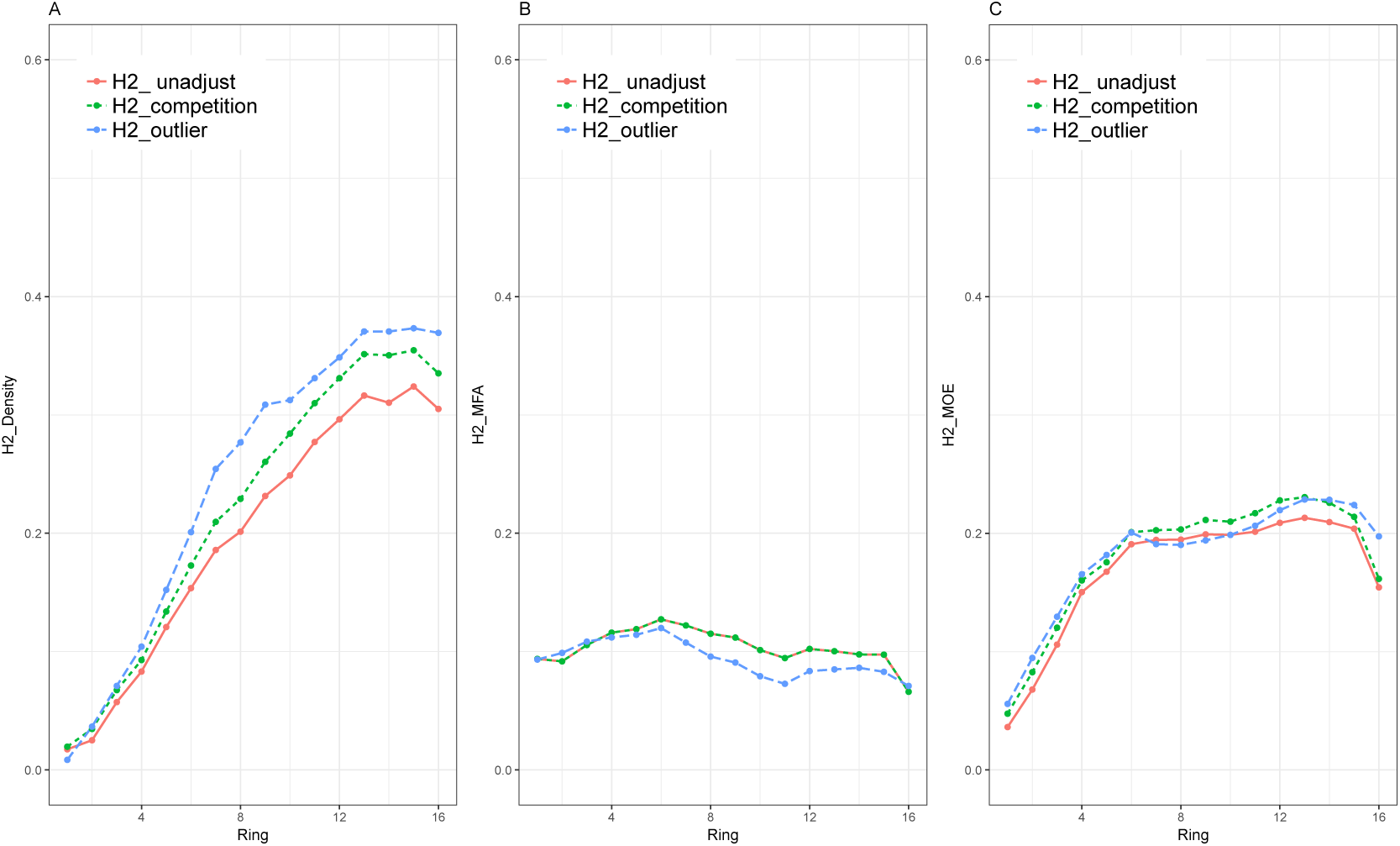
Heritability estimates using parents-offspring regression of area-weight data for area-weighted averages calculated from SilviScan data to represent the stem cross-sections at each cambial age (ring number), and using three types of data: 1) un-adjusted, 2) after competition adjustment and 3) with outliers removed for density (A), MFA (B) and MOE (C).

#### Breeding value (BV) of mothers based on progeny tests

A linear mixed model used for the estimation of parental BV and variance components can be expressed in matrix form as:

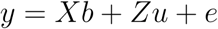

where *y* is a vector of measured data, *b* is a vector of fixed effects with its design matrix *X*, *u* is a vector of random effects with it design matrix *Z*, *e* is a vector of residuals. Fixed and random effect solutions are obtained by solving the mixed model equation [White and Hodeg, 1989]:

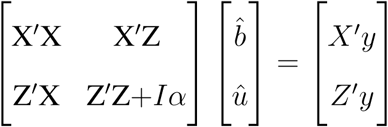

where b is the fixed effects including site, block within site and provenance, u is the random effect which is the family. I is the identity matrix with dimensions equal to the number of mothers, *α* is a ratio of residual variance and genetic variance explained by the random family effect.

### Pearson correlation

For the Silviscan data, Pearson correlation was calculated between the mother tree BVs estimated from progeny trials and the mother tree phenotypic data from the archive. For Pilodyn and Velocity, Pearson correlation was calculated between mother tree BVs estimated from progeny trials and the mean phenotypic value of mother trees. For indirect measurements the mean phenotypic value of mother trees based on data from one and three ramets, and the Silviscan data based on only one ramet sampled.

### Population heritability

As stated above, two methods for calculating heritability were compared. The first one is based on parental-offspring regression. We use linear regression to model the mother-offspring pairs for each trait value:

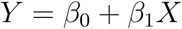

where Y is the phenotype value for the offspring and X is the phenotype value for a mother. Since genetic covariance between parents and offspring is equal to 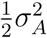 [Falconer and Mackay, 1996], we can get

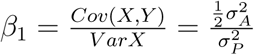

The individual tree narrow-sense heritability is

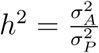

So from the slope of the regression, the estimation of the *h*^2^ can be obtained from

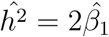

This way, the parent-offspring based heritability was computed for SilviScan data and standing tree data for Pilodyn, Hitman velocity and estimated MOE.

The second method is one based on half-sib family progeny analysis where *h*^2^=4^*^variance between families/variance of phenotype. In our study, the heritabilities were calculated for each cambial age (ring number) from the area-weighted averages representing stem cross-sections.

### Progeny size effect on heritability

In order to investigate how many progenies are needed when using parent-offspring regression for heritability estimation with multiple (one to three) ramets of the mother trees, we used a subset of progeny trees where each family had exactly six progenies in each of the two trails. Thus, in total, 180 families and 2160 progeny trees were included in the analysis. From this subset, one to six progenies were randomly selected per family, with site and heritability being estimated using parent-offspring regression. The process was bootstrapped 500 times, the means and standard errors of the heritability were then estimated for comparison.

## Results

### Phenotypic-breeding value (BV) correlation

The averages of SilviScan data across rings of same number of trees and families for the clonal archive and progeny tests, respectively, were plotted versus ring number in Figure 1 for comparison. In the progeny trials, the ring widths increased in the rings closest to the pith, reaching 5 (mm) in ring 3. Then it decreased to stablilize at 2 mm (ring 10), when wood density started to increase from a previously rather stable level. Such patterns are expected as reactions to increasing competition. Adversely, in the archive, the ring width increased and wood density decreased steadily across the first 10 rings, after which they turned to show changes in opposite directions, patterns which are expected as reactions to successively reduced competition. For MFA and MOE, the curves representing archives and trials show similar characters, respectively.

The correlations between BVs estimated from progeny and phenotypic values per ring for mother tree are presented in Figure 2. For density and MOE, correlations increased with annual rings (i.e., higher correlations at the outer most rings) for the un-adjusted, as well as, for the competition adjusted and outlier removal data. However, the last two or three annual rings experienced a decrease in correlation values. Competition adjustment and outlier removal resulted in increased correlations for density and MOE for the majority of rings. When using un-adjusted density data, the correlations increased steadily with ring number from 0.03 to 0.40, after competition adjustment the maximum correlation reached 0.43, and after outlier removal it reached 0.47. In the case of MOE, correlations using un-adjusted data increased from 0.06 to 0.25 at ring 6. It continued to increase marginally to 0.26 at ring 14, after which it decreased in the outermost rings. Competition adjustment and outlier removal followed similar patterns and reached maximum correlations of 0.27 and 0.28, respectively. In the case of MFA, correlations decrease after annual ring 6. Both adjustment and outlier removal caused correlation decrease for all rings. Correlations for the un-adjusted MFA data ranged from 0.18 reached at ring 6 to 0.05 in the outermost ring. After competition adjustment and outlier removal the highest correlation reached were 0.17 and 0.16, respectively. While the lowest correlation dropped to minimum values of 0.02 and 0.05,.

The estimated correlations between BV of progeny and phenotypic value of mother tree were 0.29, 0.13 and 0.23 for Pilodyn, Velocity and MOE(ind), respectively. When using three ramets of the mother trees, the correlation increased to 0.32, 0.15 and 0.28, respectively.

### Heritability

Heritability estimations for stem cross-sections at each age (ring number) are presented in Figure 3 on SilviScan data and one single ramet from the archive. For density, they increase steadily from low level up to 0.32 at age 15 years. Whilst when using un-adjusted data, it increased to 0.36 and 0.37 with the competition adjusted and outlier removed data, respectively. For MOE all types of data provide similar graphs, increase from low level to roughly ring 6, then marginal increases to 0.21 for un-adjusted data and 0.23 for the two adjusted types at ring 15. For MFA, heritabilities using all types of data ranged across all rings between 0.07 and 0.13 with maximum values of 0.13 at ring 6 for all alternatives. This was followed by a slow decline towards the same minimum value at ring 15, during which the outlier removed graph showed a somewhat lower path. Thus, the data adjustments had clear positive effects on heritability only for wood density, with 13% for competition adjustment and 16% for outlier removal from a level of 0.32. For MOE they gave slight improvements from the level of about 0.20, while no benefits were obtained from these types of data adjustments for MFA.

Heritability estimations of the whole stem cross-sections based on progeny data and parent-offspring regression are presented in Table 1. Heritability estimates based on parent-offspring regression were higher for density (0.35), MFA (0.15) and MOE (0.28) measured with Silviscan compared to use of the indirect methods and phenotyping of only one ramet from the archive: Pilodyn (0.27), Velocity (0.13) and MOE(ind) (0.13), although not significantly if considering the standard errors. However, when indirect measurement data from three ramets of the parent were analysed they gave higher estimates for all three wood quality traits (0.41, 0.19 and 0.30) than use of SilviScan data and only one ramet. Heritability estimates based on half-sib progeny correlation were higher for all three wood properties (0.43, 0.29, 0.38) than parent-offspring heritability based on a single ramet data using Silviscan, and also higher than indirect measurement methods (0.31, 0.20, 0.28) in half-sib progeny. However, heritability estimates based on half-sib progeny correlation were similar to the heritability estimates based on the parent-offspring regression of indirect method using three ramets of the parent for all three wood properties.

**Table 1:** Heritability and repeatability estimates based on measurements of wood density, MFA and MOE with SilviScan from increment cores: area-weighted averages representing the cross-section of the stem at different cambial ages (ring numbers), density, MFA, MOE), and based on indirect measurements on standing trees of their proxies with Pilodyn and Hitman (velocity), from which also MOE(ind) was estimated. For comparison, all these heritability estimates were based only on the 162 families for which data on three ramets from the archive is available.

| Methods | SilviScan |  |  | Indirect methods |  |  |
| --- | --- | --- | --- | --- | --- | --- |
|  | Density | MFA | MOE | Pilodyn | Velocity | MOE<sub>ind</sub> |
| Parent-offspring regression_1 ramet | 0.35 ($\pm 0.16$) | 0.15($\pm 0.16$) | 0.28($\pm 0.16$) | 0.27 ($\pm 0.15$) | 0.13 ($\pm 0.15$) | 0.13 ($\pm 0.15$) |
| Parent-offspring regression_3ramets | N/A | N/A | N/A | 0.41 ($\pm 0.15$) | 0.29 ($\pm 0.15$) | 0.30 ($\pm 0.15$) |
| Half-sib correlation (offspring only) | 0.43 ($\pm 0.09$) | 0.29 ($\pm 0.08$) | 0.38 ($\pm 0.08$) | 0.31 ($\pm 0.08$) | 0.20 ($\pm 0.08$) | 0.28 ($\pm 0.08$) |
| Repeatability | N/A | N/A | N/A | 0.52 ($\pm 0.06$) | 0.30 ($\pm 0.04$) | 0.45 ($\pm 0.05$) |

The repeatabilities of Pilodyn, Velocity and MOE(ind) of mother trees were 0.52, 0.30 and 0.45, respectively, which were higher than progeny-offspring and half-sib progeny heritability estimates based on both Silviscan traits and indirect traits

### Effect of progeny size on parent-offspring heritability estimate and standard error

The most prominent consequence of increasing the progeny size is the decrease in the standard errors (i.e., more precise estimation of heritability) (Figure 4). When a progeny size four was selected, heritability stabilizes for MOE, whereas it decreases for Velocity and reaches a maximum value at progeny size 6 for the Pilodyn trait. The mean heritabilities of Velocity, Pilodyn and MOE(ind) are in the range of 0.26 to 0.28, 0.56 to 0.59 and 0.32 to 0.34 respectively.

**Figure 4:**
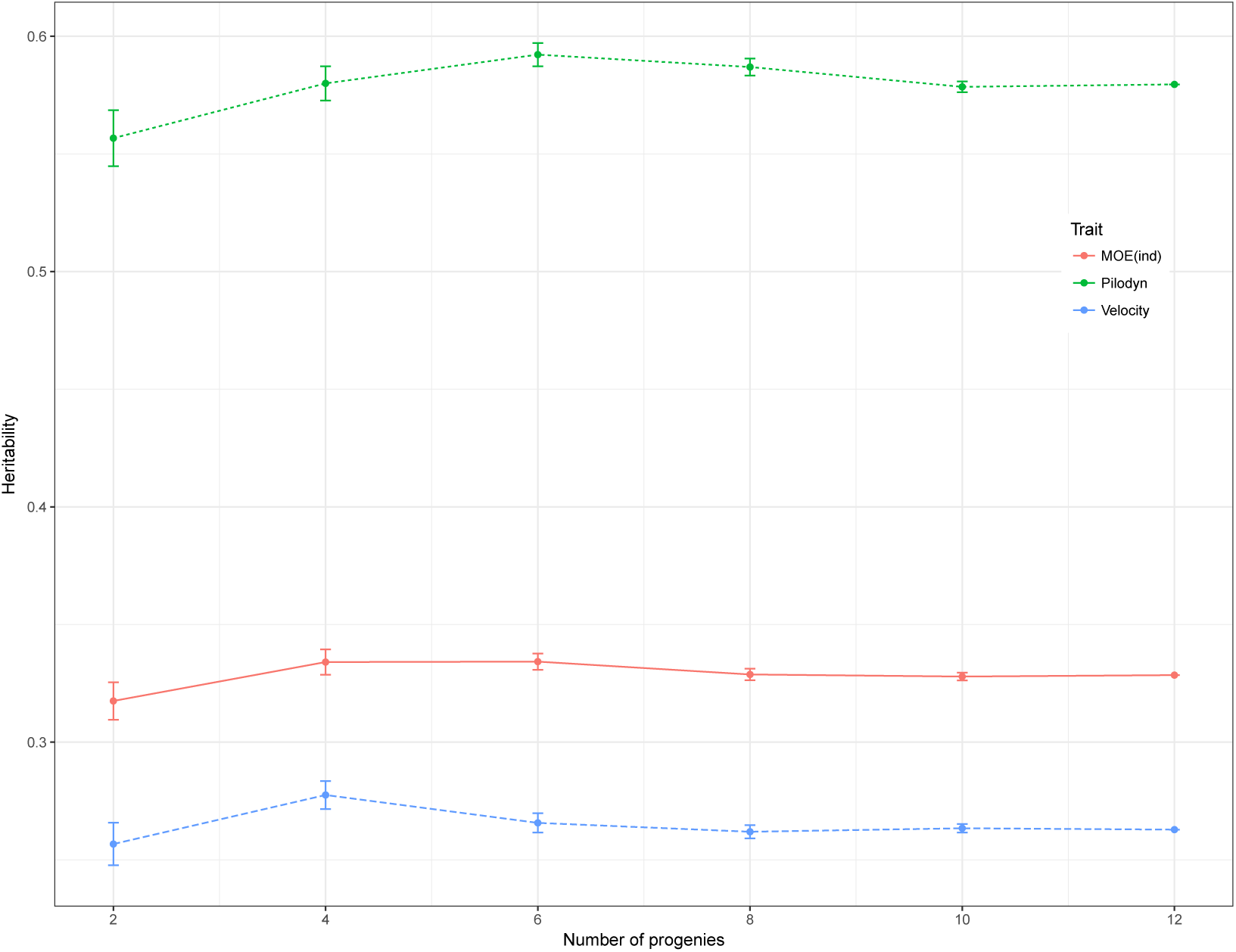
Heritability estimation based on different number of progenies for each mother and estimated by parent-offspring regression for Pilodyn, Velocity and MOE(ind). The number of ramets per mother clone varied among plus trees from one to three.

## Discussion

### Low to moderate correlation between clonal archives and progeny tests

In operational breeding, selection of clones for crossings to the next generation (backward selection) is usually conducted through evaluation of their half-sib or full-sib progenies grown in common-garden experiments, so called progeny trials. This is a method that involves multiple actions such as crossing (in case of full-sib progenies), seedling production, seedling establishment (often in multiple sites), and assessment and evaluation of multiple tree properties. The high demand in time and costs of such a process motivated the first question of this study: could the mothers be ranked based on tree properties assessment directly on the trees of the mother clone/genotype from the clonal archives? To answer this question, we estimated solid wood properties, known to be highly heritable and little G x E, in 524 mother clones in a clonal archive and in their half-sib progenies established in two test sites.

Our study reveals a low to moderate correlation between the phenotypic values assessed directly in the mother trees and the breeding values (BV) estimated on their half-sib progenies. This result is similar with use of both the Silviscan and the indirect measurements. Among the three solid wood properties, wood density had a moderate correlation between parent and progeny except at young ages, and unstable at the highest cambial age. The correlation reached above 0.4, on use of un-adjusted data at cambial age 13, for competition adjusted data at ring 11, and when outliuers were removed at ring 7. MFA/Acoustic Velocity had the lowest correlation, with no age reaching higher than 0.19. MOE showed intermediate correlation, reaching above 0.2 after age 4, and for this trait, adjustment by outlier removal or competition index brought little improvement to the estimates. Despite the high heritability of theses solid wood properties [Hannrup et al., 2004, Steffenrem et al., 2009], it cannot be ruled out that the management activities, such as grafting, topping and thinning, may have affected the correlation between mother parent and progenies. However, the low and moderate correlation between parent and progenies did not affect greatly the estimates of heritabilities compared to estimates from progenies only.

The heritability estimates for density, MFA and MOE obtained in our study, using the tested approach based on SilviScan data from only one single ramet from the archive was lower than previous reported for wood density and MFA in white spruce (0.65, 0.36 and 0.29; [Lamara et al., 2016], Scots pine (0.40, 0.39 and 0.30; [Hong et al., 2015] and lodgepole pine (0.48, 0.31, 0.21; [Hayatgheibi et al., 2017]. If the SilviScan data was replaced with data from the indirect methods, the heritabilities decreased. However, if use of the indirect methods was combined with an increase of up to three ramets from the archive, the heritabilities of pilodyn was higher and comparable with such reported for Scots pine, but lower than the values observed for white spruce and lodgepole pine. Under these conditions, the heritability for MOE(ind), was close to that for white spruce and Scots pine, but higher than the values for lodgepole pine. For velocity, the heritability was lower than in all those previous studies. Thus, it is reasonable to believe that the heritability had been higher and become comparable with other species if we had SilvScan data for more ramets. We also estimated relatively high clonal repeatability for Pilodyn (0.52) and MOE(ind) (0.45), which are higher than the narrow sense heritability. Both non-additive genetic variation and imperfect experimental design in the archive may have contributed to such higher clonal heritability.

### Heritability estimates improve with number of ramets and progeny size

In conifers, narrow-sense heritability is often based on half-sib progeny correlations. As an alternative, narrow-sense heritability can be computed from parent-offspring regression [Falconer and Mackay, 1996]. This approach has been commonly used in crops [Casler, 1982, Fernandez and J. C. Miller, 1985, Vogel et al., 1980]. More recently, it has also been applied for wood quality traits (wood density and spiral grain) in Norway spruce [Steffenrem et al., 2016]. The author reported similar heritabilities and standard errors for both methods of narrow-sense heritability computing, parent-offspring regression and progeny data. This is in line with results mentioned in simulation studies on wild populations of both binary and continuous traits [de Villemereuil et al., 2013], and on real data using eight morphological traits of a natural population of the great reed warbler (Acrocephalus arundinaceus) [kesson et al., 2008]. In our study, Silviscan-based narrow-sense heritabilities based on progeny were higher, but comparable to results based on parent-progeny regression using the indirect method, where data for all three ramets of the mother trees was available. Thus, we can conclude that the highest values of narrow-sense heritability estimates were obtained for the parent-offspring regression, as far as data from at least three ramets is involved. Therefore, in agreement with Steffen-rem et al (2016), also we conclude that parent-offspring regression results in higher values of heritability. We observed that increasing progeny size increases precision of the heritability estimates (i.e., lower standard errors). Moreover, increasing progeny size has its major impact in the transition from two to four progenies, whereas further increases do not result in a clear increase in the heritability values.

## Conclusion

Our study resulted in the following conclusions:

- The correlations between the phenotypic values of the mother trees and the breeding values estimated on their half-sib progenies are low to moderate. Reasons for this may be management practices and lack of experimental design in archives, contributing to the parent and progeny correlation.
- Offspring progeny heritability estimates based on SilviScan measurements were higher than parent-offspring regression using a single ramet from the archive. Both these approaches resulted in higher heritabilities than when traits had been determined with more efficient indirect measurements using Pilodyn and acoustic Hitman on the standing trees.
- When three ramets per plus tree were measured using Pilodyn and Hitman, the parent-offspring regression heritability estimates were higher than those based solely on progeny data.
- The influence of progeny size used was studied for the latter approach. The standard error of the heritability estimates decreased with increasing progeny size, and stabilised at four progenies for the heritabilities of all traits investigated. Parent-offspring regression could thus be used to estimate heritabilities using clonal archive parent for the investigated solid wood properties. Three ramets and four progenies of each parent were sufficient for reaching the same heritability levels as with more extensive analyses of progeny trials.

## Acknowledgments

We acknowledge Skogforsk for support on the collection of data in both the clonal archive and progeny trials, and also Åke Hansson, Thomas Trost and Fredrik Adaås, Innventia, now RISE Bioeconomy, for the excellent work with the Silviscan wood analyses. We also acknowledge Bio4Energy and the Swedish Foundation for Strategic Research (SSF, grant number RBP14-0040) for support to conduct this study.

